# Long-Term Imprint of Prenatal War Trauma on Brain Structure: Evidence from Genetically Informed Neuroimaging

**DOI:** 10.1101/2025.11.05.686844

**Authors:** G. Aydogan, S. Kollmorgen, V. Mante, R. Karlsson Linnér, B. Kleim, G. Nave, C.C. Ruff

## Abstract

The global rise in armed conflicts has raised concerns about their long-term consequences. While the immediate mental and physical impacts of war are well documented, the effects on unborn children of expecting mothers remain poorly understood. Here, we leverage a large dataset (*N*=37,856) combining historical World War II bombing records with brain imaging and genetic data to examine whether in-utero exposure to air raids is associated with lasting differences in adult brain structure, cognition, and physical health. Using a pre-registered quasi-experimental design controlling for genetic and environmental factors, we find that exposure during the last trimester within 2 km of bombings is associated with lower gray matter volume, which partially mediates adverse effects on intelligence and obesity. These findings suggest a plausible pathway linking prenatal trauma to long-term differences in neurodevelopmental and functional outcomes, underscoring the vulnerability of the fetal brain to acute stress and highlighting the importance of protecting pregnant women during armed conflicts.

## Main Text

War stands out as one of the most severe yet avoidable tragedies in human history(*1–4)*. The persistence of state-based armed conflicts since the end of World War II (WWII) has had profound effects on global stability, resulting in the displacement of millions of people(*5)* with severe personal and economic repercussions(*1)*. In 2023, despite the absence of a global conflict on the scale of WWII, the global economic impact of armed conflicts was estimated at $19.1 trillion, equivalent to 13.5% of global GDP(*6)*. Beyond its economic toll, the psychological effects of warfare continue to harm both soldiers(*7, 8)* and civilians alike(*9, 10)*, with long-lasting mental health consequences including diminished cognitive abilities(*11)*, depression(*12)*, and post-traumatic stress disorder(*12, 13)*.

While the psychological burden of war trauma is well documented, the cross-generational effects of war remain less understood(*14)*, specifically on unborn children of expectant mothers who are likely to experience heightened stress during such times of extreme adversity. Most causal evidence about the effects of maternal stress on unborn children stems from animal studies, which have illuminated molecular and neurobiological pathways and mechanisms through which prenatal stress can alter fetal development. These pathways include, for example, inhibited N-methyl-D-aspartate (NMDA) receptor gene expression in the hippocampus and frontal cortex(*15, 16)*, reduced myelination in the corpus callosum(*17)*, and suppressed neurogenesis of hippocampal granule neurons in the developing fetus(*18)*. Further, high levels of the key stress hormone cortisol during pregnancy have been shown to impair proliferation and neuronal differentiation in the hippocampus of the developing nervous system(*19)*. However, evidence generalizing these effects to human populations is scarce, and it remains unclear whether these observed effects would translate to effects of war trauma on human gestation at all.

Only a few small prospective studies in humans have investigated the effects of prenatal stress on neuronal development (*20)*, but this has rarely been done in the context of war. Nevertheless, exposure to non-war-related prenatal stress was found to be negatively associated with cortical thickness primarily in frontal and temporal brain regions of the developing fetus(*21, 22)*. This reduction in cortical thickness, and other gray matter alterations, were associated with decreased learning and memory(*23, 24)* as well as disrupted emotional regulation(*22, 25)*, impaired cognitive performance(*23, 26)*, and increased vulnerability to mental disorders later in life(*27, 28)*. Similarly, higher prevalence of depression, anxiety, and related psychiatric disorders in adulthood was also linked to prenatal stress(*29–33)*. Such effects may in principle be most marked for war-related prenatal stress, which represents a uniquely harmful condition due to its chronic, pervasive, and multifaceted nature(*14)*. Unlike isolated traumatic events such as accidents or personal loss, war imposes a sustained psychological and physiological burden through ongoing threats to safety, displacement, disrupted healthcare, and loss of social support. This prolonged stress can lead to persistently elevated maternal cortisol levels(*14, 28)*, which cross the placental barrier and may interfere with fetal brain development. Consistent with this, data from the Korean War show that individuals exposed to in-utero trauma during the most intense phases of conflict had significantly poorer socioeconomic and health outcomes later in life, including lower educational attainment and higher disability rates(*14)*.

Despite these suggestive findings, causal evidence of the impact of war on neural development remains elusive, for two main reasons: First, due to obvious ethical and practical considerations, no randomized controlled studies have investigated the causal relationship between warfare and alterations in brain development. Thus, even though correlational studies suggest a plausible link between prenatal trauma and altered neural development(*34)*, these links may not reflect causal relationships but rather confounding genetic or environmental factors that influence both the occurrence of trauma and neural development(*35, 36)*. Second, while the possibly small effect sizes of trauma on brain development would require studies with large samples(*35–37)*, the high costs and limited data availability have restricted previous neuroimaging research to relatively small sample sizes, usually on the order of less than hundred participants(*21, 38, 39)*. Previous work has therefore been underpowered to detect the corresponding effects and estimate their magnitude precisely(*40)*. Finally, the existing small studies could not systematically control for potentially confounding genetic and environmental factors(*35, 36)*.

Here, we examine whether in-utero war trauma leaves a lasting imprint on brain structure and subsequent life outcomes decades later. To this end, we leverage a unique natural experimental setting of war-induced prenatal trauma within a rich, large-scale brain imaging dataset that includes environmental and genetic controls. Specifically, we combine the largest collection of publicly available brain images from *N* = 37,856 participants in the UK Biobank (UKB)—of which 6,665 participants were born during WWII (see Fig. 1A and Fig. S3)(*41)*—with data from the British National Archive that include meticulously recorded bombing locations from German air raids, detailing 32,870 air raid locations in the UK between 1939 and 1945 (see Fig. 1B)(*42)*. Using the combined dataset, we investigate in a pre-registered study how prenatal trauma of expectant mothers during WWII, via exposure to air raids within different degrees of proximity to birth location and different trimesters of pregnancy, affected the brain development of their unborn children, while systematically controlling for genetic and environmental factors. While this study uses a quasi-experimental design and extensive controls for confounding factors, we cannot fully rule out all potential sources of unmeasured confounding. Therefore, to mitigate those concerns, we perform extensive robustness and sensitivity analyses to probe our findings and to assess the plausible link between in-utero war trauma and its lasting imprint on the brains of those exposed.

**Fig. 1:**
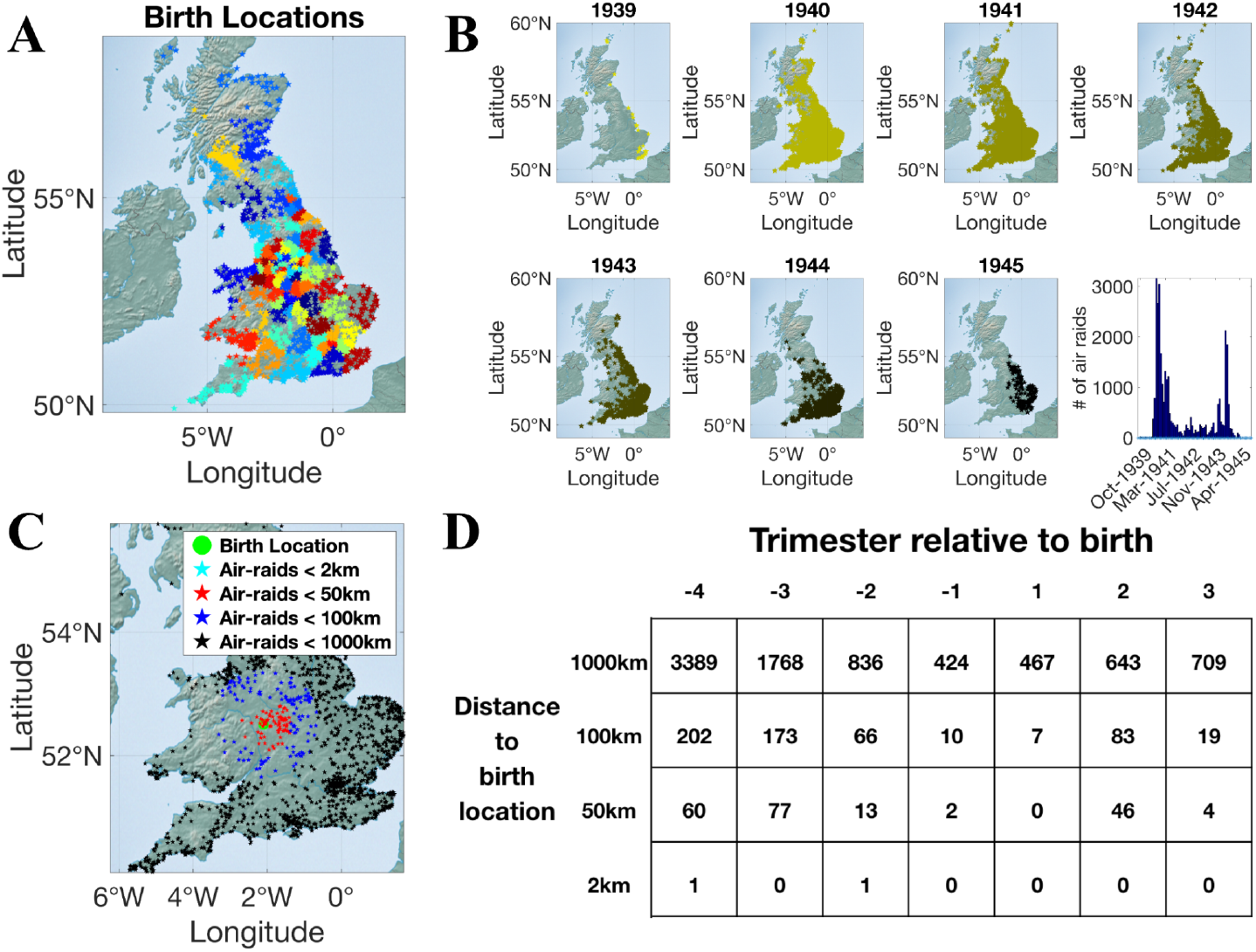
(A) Geographical birth locations of all participants (including those with no MR scans), binned into 100 clusters. These were used as dummy variables and to cluster standard errors. Each star represents a participant’s birthplace (non-jittered), while the colors indicate 100 geographical clusters, generated using a *k*-means clustering algorithm with *k*=100 and 10,000 iterations following random seeding. (B) Bombing locations over time within the UK. While during the first bombing wave in 1940 (known as the Blitz) virtually all populated areas in the UK were exposed to bombings, there is substantial variation over time, rendering a systematic incorporation of spatial and temporal controls necessary. (C) To illustrate our identification strategy, we plotted the birth location of one example individual and the bombing locations within 9 months before the person’s birth. (D) Illustrated is the number of air raids that this person was exposed to in utero as a function of spatial (2km, 50km, 100km, and 1000km) and temporal (ranging from 12 months before to 9 months after birth, binned in trimesters) distance to the person’s birth. The table illustrates the substantial variation in the number of air raids over time (and distance) relative to this individual’s birth.

### Identifying effects of in-utero trauma associated with air raids

To support a causal interpretation of the link between air raids and brain development, air raid locations must vary randomly across time and space, precluding systematic differences in strategic avoidance of exposure between participants. Our analysis leveraged the historical context of Nazi Germany’s World War II (WWII) terror bombing campaign, which exposed the British civilian population to widespread and largely unpredictable aerial attacks, irrespective of individual characteristics or locations(*43)* (see Fig. 1B). These air raids were highly indiscriminate compared to modern precision bombing, due to factors like evasive maneuvers, night-time missions, and pilot errors, resulting in targeting errors that often spanned many kilometers(*43, 44)*. Further, the Germans’ goal was to terrorize the British population indiscriminately, providing a disincentive to bomb specific locations at predictable times, even if this had been possible. This near-randomness in air raid exposure serves as the basis for our quasi-experimental setting (see Methods for details). Furthermore, recognizing that certain areas (e.g., large cities) were still bombed more frequently than others (e.g., rural areas), our analyses account for correlations within geographical areas by controlling for (and clustering standard errors on) 100 geographical regions across the UK derived from naturally occurring clusters of birthplaces. We identified these clusters through a data-driven algorithm (*k*-means), which groups individuals born in geographically similar areas to avoid potential bias from arbitrary or manually defined boundaries (see Fig. 1A and Methods for details). This allows us to control for general area-specific effects while effectively comparing mothers within the same geographic region over time, contrasting exposure to air raids within different geographical radii and at different time points from conception to birth. For example, expectant mothers who gave birth right before a major air raid (placebo) can serve as controls for mothers in the same geographical area who were exposed to the same air raids, but within their last trimester of pregnancy (treatment).

To identify how the effect of prenatal exposure to air raids depends on their temporal and spatial distance to birth, we binned the number of exposed air raids using four spatial radii, from immediate life-threatening proximity (2km) to distant areas unaffected by raids (1,000km) (see specification-curve analysis and Fig. S1 in SI), and across seven temporal windows ranging from before conception to pregnancy and finally to 9 months after birth (see Fig. 1C-D). The analyses using different spatial distances allowed us to quantify effects related to varying war experiences, ranging from hearing sirens or distant bombings without immediate consequences to more immediate, consequential experiences, such as the ground shaking from nearby blasts and destruction and death in the immediate surroundings. To account for the uncertainty in self-reported birth or air raid locations, and differences in the number of participants we could include in each cell of the analysis, we conducted a specification-curve analysis to assess the stability with which different radii in close proximity to the bombings (<10 km) yielded consistent effects on GMV (see Fig. S1 in SI). This analysis indicated that a 2-km radius was the most robust to positional granularity. Accordingly, we use a 2-km radius in all subsequent analyses. Crucially, our analyses systematically controlled for various genetic and environmental factors known to correlate with Gray Matter Volume (GMV)(*36)*. Specifically, we included 40 genetic principal components (accounting for latent population structure), SES proxied by current residence, birth year dummies, genetic sex, age at scan, handedness, volumetric scaling of the T1 image, as well as Total Intracranial Volume (TIV) into our analyses (see Methods).

### In-Utero Air Raids are Associated with Gray and White Matter Volume

To investigate whether prenatal war trauma has lasting effects on brain development, we examined GMV and white matter fiber structure, two well-established neuroimaging markers of neuronal integrity and maturation. While GMV reflects the density and size of neuronal cell bodies(31), dendrites(32), and unmyelinated axons(33) in specific brain regions, white matter fiber integrity captures the organization and connectivity of axonal pathways that link distant brain regions(*45–47)*. Both measures have been associated with key cognitive functions, including memory(*48, 49)*, processing speed(*45)*, and executive control(*35, 50)*, as well as emotion regulation(*50)* and mental health outcomes(*47, 51)*. Moreover, GMV reductions have been found in children exposed to in-utero stress(23), which is thought to reflect disruptions in key neurodevelopmental processes, such as neurogenesis, synaptic pruning, and neuronal loss(22).

First, we tested whether exposure to air raids—specifically, the number of air raids within close temporal and spatial proximity to birth—affected the total GMV of exposed participants. To achieve this, we first regressed total GMV on the number of air raids of all 28 distinct space-time bins (see Fig. 1D) while including all standard control variables to account for environmental and genetic confounds. This revealed that air raid frequency within close proximity (less than 2km) to the birth location exhibited a significant association with total GMV measured later in life, specifically when air raids occurred within the last trimester of pregnancy (standardized *β* = −.011; 95% CI −.016, −.007; *t*(6,090) = −5.38; *P*<7.7×10^−8^, with 100 clustered standard errors, two-sided) (see Fig. 2A). Bombings at greater spatial distances, or in different trimesters of pregnancy (see Fig. 2A), had no discernible effect on GMV (*Ps* > .01).

**Fig. 2:**
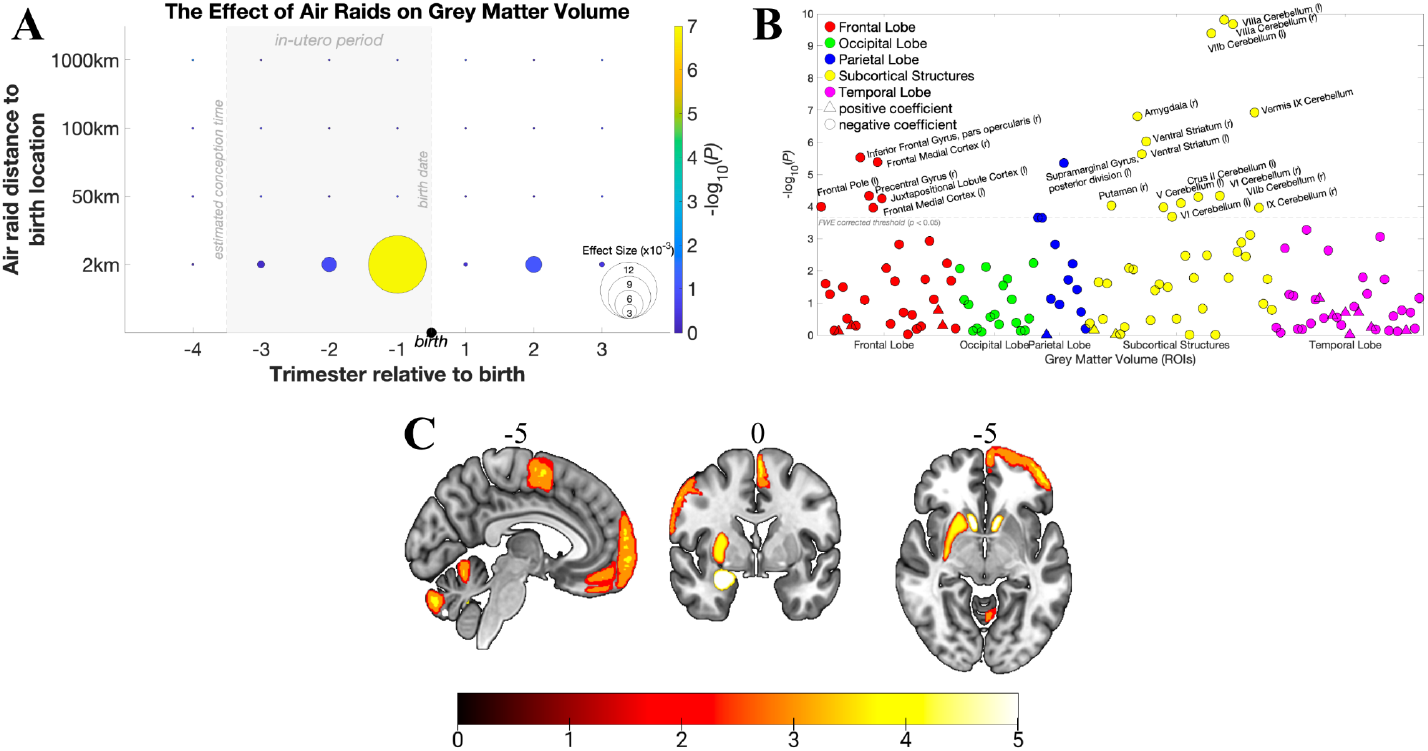
(A) A heat map of *p*-values of a regression model estimating the effect of air raids on total GMV, controlling for all standard and genetic controls. The size and the color of the circle indicate the size of the effect and the related negative log transformed *p* values, respectively. Bombings within 2km radius of birth location within the last trimester show a significant negative effect on GMV (*P*< 7.7×10^-8^). In contrast, air raids with greater temporal or spatial distances to birth show no significant association. Panels (B) and (C) illustrate the negative log-scaled *p*-values of ROI-based analyses, which indicate that in-utero exposure to air raids were related to localized effects, mainly in frontal and cerebellar brain structures (*P*_*FWE*_ < .05).

This specific association of GMV with bombings in close proximity during the last trimester was robust to analyses considering the whole data set, i.e., when including participants born before or after the war who were not exposed to any bombings in utero (standardized *β* = −.011; 95% CI −.015, −.007; *t*(37,550) = −5.74; *P*<9.5×10^−9^, with 100 clustered standard errors, two-sided). This further supports that the effects of in utero bombings are robust to the included control variables (such as age, location, genetic envelope etc.), since these are estimated with higher precision in the full dataset. Next, we investigated potentially localized effects of prenatal trauma in the brain, as some brain areas might be more susceptible to traumatic events during pregnancy (see Fig. 2B-C). To this end, we regressed the GMV of 139 regions (defined a priori based on the Harvard-Oxford atlases provided by UKB) on the number of air raids (across all trimesters and distances), again including all standard controls. To account for multiple hypothesis testing, we applied a family-wise error rate of *P*_*FWE*_ = .05, estimated based on a permutation test (1,000 permutations). Consistent with suggestive findings in much smaller datasets (*21, 23)*, our analysis identified 21 significant regions (within 2km radius and last trimester bin) primarily in frontal areas such as the left Frontal Pole (l), bilateral Medial Prefrontal Cortex (bilaterally), the right Inferior Frontal Gyrus (r), and regions of the cerebellum (*P*_*FWE*_ < .05), with standardized *βs* ranging from −.029 and −.055 (see Fig. 2B-C as well as Table S1). We observed no significant effects in the temporal lobe, despite previous reports of such effects.

Alongside the GMV alterations, we also found significant effects (*P*_*FWE*_ < .05, standardized *βs* ranging from −.073 and .054) on various white matter fiber tracts (29 out of 432) (see Fig. S4 and Table S2), as indicated by different fiber integrity metrics such as lower fractional anisotropy in the genu of corpus callosum and lower diffusion tensor mode in the superior fronto-occipital fasciculus and in the posterior thalamic radiation, which are white matter structures related to cognitive abilities such as short-term memory and fluid intelligence (IQ)(*52–54)*. Notably, most of the GMV and WM structural changes were concentrated in brain regions involved in higher-order cognitive processing and executive function, pointing to a possible neurobiological pathway by which in-utero war exposure may be associated with adverse long-term effects, such as impaired cognitive abilities, poorer health outcomes, and reduced employment status(*14)*.

### Trauma-Related Gray Matter Alterations and Real-Life Outcomes

To examine whether war-trauma-related GMV reductions might have long-term downstream effects on cognition and health(*14)*, we tested whether reduced GMV mediates the link between prenatal air raid exposure and later-life outcomes. Specifically, we examined whether individuals with lower GMV following prenatal trauma also showed changes in IQ and BMI (body mass index) in adulthood, while again controlling for genetic and other confounds (see Fig. 3A-B). The mediation analysis revealed that lower GMV in our sample was associated with lower IQ (standardized *β* = .096; 95% CI .05, .14; *t*(5,295) = 3.94; *P*<8.2×10^−5^; with 100 clustered standard errors, two-sided) and higher BMI (standardized *β* = −.2; 95% CI −.*25*, −.*15*; *t*(6,082) = −7.68; *P*<1.8×10^−14^, with 100 clustered standard errors, two-sided). While the direct associations between bombings and IQ or BMI were not statistically different from zero (*Ps* > .01), the indirect effects were highly significant (*Ps* < .001), indicating that total GMV reductions statistically mediated the association between bombings (within 2km radius and in the last trimester) and IQ or BMI. Such mediation pattern (*55, 56)* may be consistent with a vulnerability-based mechanism (*57–59)* - that is, the cognitive consequences of prenatal trauma appear not to be driven by direct exposure in every person, but rather seem to emerge via their impact on neurodevelopment. These findings highlight GMV as a critical neurobiological mediator of potential long-term cognitive consequences of prenatal trauma. Thus, GMV reductions may represent a plausible pathway linking prenatal trauma to enduring cognitive outcomes, consistent with associations reported in prior correlative studies(*21, 23)*.

**Fig. 3:**
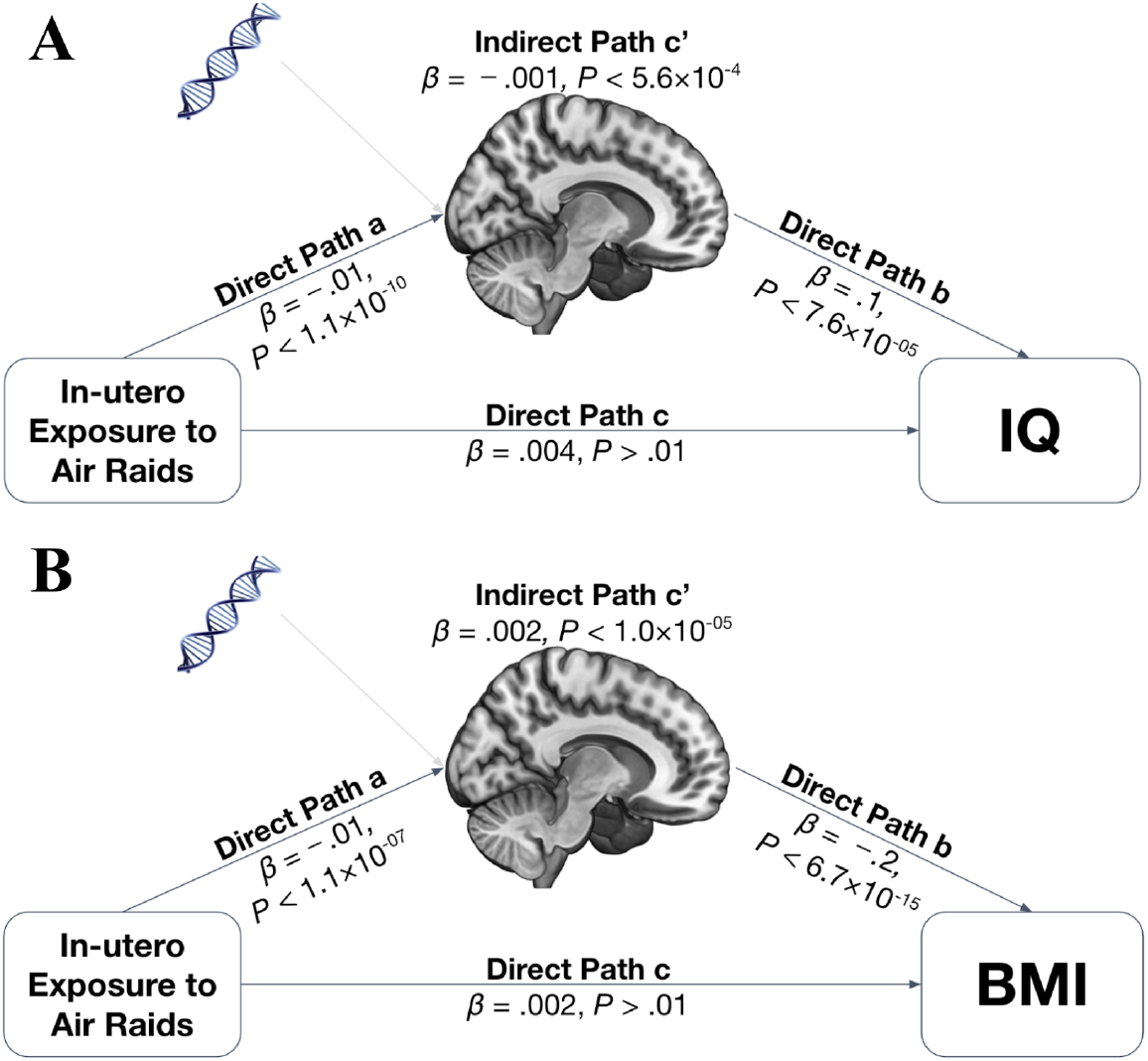
(A) Mediation analysis showing that the association between air raid exposure (last trimester within 2km radius) and IQ is significantly mediated by differences in total GMV (*P*<.001). (B) Similarly, the effect of air raid exposure (last trimester within 2km radius) on BMI is also significantly mediated by differences in GMV (*P*<.001). Both mediation analyses exhibit significant indirect effects in the absence of direct effects, indicating that the pathway through GMV reductions is critical for observing these cognitive and physical health outcomes of air raid exposure.

### Addressing Potential Confounds and Selection Biases

Previous research suggests that prenatal stress due to air raids and ensuing elevated cortisol levels might lead to premature births or even miscarriages, which might potentially bias our findings, as low birth weight and preterm birth have previously been related to reductions in regional brain volumes(*60)*. To ensure that higher rates of premature births do not confound our results, we first regressed birth weight—a reliable proxy for premature birth(*61)*—on air raid exposure and found no significant association (*Ps* > .05, *N* = 3,180), indicating that premature births did not significantly increase due to the war. Moreover, the absence of any effect of postnatal trauma on GMV further suggests that environmental exposures such as damaged infrastructure, pollution, or physical injury are unlikely to explain the observed associations. Together, these findings support the interpretation that the effects of air raid exposure on brain development are specific to the prenatal period and may not be driven by general environmental adversity.

To investigate the possibility that higher miscarriage rates may have influenced our results, we consulted population-level statistics from the British Office for National Statistics (ONS). This confirmed that miscarriage rates did not rise during the war - in fact, stillbirth rates in England and Wales significantly declined from 38.3 stillbirths per 1,000 births in 1938 to 27.6 by 1945, a reduction of nearly 40%(*62)*. Due to these data it appears unlikely that miscarriage rates confounded our results.

To account for the possibility that ‘smarter’ expectant mothers might better shield themselves from the effects of bombings—through avoidance or precautionary measures like seeking shelter earlier—we used each participant’s polygenic risk score of IQ as a proxy for inherited IQ from their parents (see SI for more details). We incorporated this proxy for parents’ IQ as a control variable in our analysis and observed no meaningful change in the effect of air raids on total GMV (standardized *β* = −.012; 95% CI −.018, −.006; *t*(5,927) = −4.04; *P*<5.4×10^−5^, with 100 clustered standard errors, two-sided), indicating that higher IQ of expectant mothers did not confound the observed effects on fetal development.

We then explored whether temporal self-selection via prospective family planning might have biased our results (e.g., if some women delayed family plans as a consequence of Nazi Germany’s terror bombing campaign). Arguably, the sudden onset of Nazi Germany’s very first terror bombing campaign (called the ‘Blitz’, ranging from September 1940 to mid-1941) can be considered largely unexpected by the general population and allows us to investigate the effect of prenatal trauma largely independent of maternal planning. By focusing on this time frame (i.e., births up to 1941), and comparing birth rates to those after 1941, we can evaluate whether the apparent effect of the air raids on brain development might be confounded by selected women’s deliberate family planning in anticipation of increased conflict. The results of this analysis mirrored those observed across the entire dataset (standardized *β* = −.013; 95% CI −.019, −.007; *t*(1,656) = −4.19; *P*<3×10^−5^, with 100 clustered standard errors, two-sided), indicating that the effects were similarly present during the initial wave of bombings and later phases. Furthermore, fertility rates from 1939 to 1945 rose by approximately 12% compared to pre-war rates, suggesting that family planning remained largely unaffected by the war(*62)*. This indicates a robust association of prenatal trauma with GMV, even in a time frame where anticipatory actions by expectant mothers, based on the predictability of conflict, can be largely excluded.

Finally, individuals exposed in utero to bombings may have had lower life expectancy, as previous reports suggest elevated mortality rates in this group(*14)*. However, if true, such attrition would bias our findings downward by omitting the most severely affected cases, thus underestimating the full impact of prenatal trauma. Similarly, some air raids may be missing from our dataset. As long as this omission is random, it would reduce measured exposure levels,again likely not inflating our results but instead underestimating the true effect. Therefore, while our conclusions remain robust, the actual impact of prenatal trauma may differ from that reported here. While residual confounding cannot be fully excluded, the pattern and robustness of our results are consistent with the interpretation that severe prenatal war trauma may leave a lasting imprint on the developing human brain.

## Conclusion

We leverage a natural quasi-experiment created by World War II air raids to study whether acute, exogenous prenatal stress is associated with altered human neurodevelopment in adulthood, using a dataset orders of magnitude larger than prior work on this topic. Exposure mattered only at very close range: prenatal stress was only associated with GMV differences when bombings occurred in very close proximity (<2km), suggesting that the intense terror and the immediate fear of life-threatening danger might be necessary to trigger such adverse effects(*38)*. Moreover, this association was virtually exclusively confined to the last trimester (28 to 40 weeks)(*39)*. This spatiotemporal specificity could stem from two main reasons: First, maternal stress may primarily impact the central nervous system of the fetus during the last trimester of gestation, which is a critical period marked by significant brain development, such as the maturation of cortical layers and substantial increases in brain volume and neural connectivity(*63)*. For instance, gliogenesis, which begins around the 22^nd^ gestational week, results in the production of region- and subtype-specific glia. And by approximately the 32^nd^ gestational week, these glial cells begin to assist in the myelination of neurons(*63)*. A disruption of this process would account for the observed reductions of GMV in frontal brain areas, which mature relatively late, as well as for the reduced white matter integrity of the corpus callosum and other critical fiber tracts. These findings closely dovetail with research on sensitive periods in brain development, which suggests that the timing and nature of trauma can shape distinct neurodevelopmental outcomes (*32, 64, 65)*. Second, the effect being limited to the last trimester may also indicate that expecting mothers may have been more likely to remain near their final birth locations during this phase. This could potentially obscure any impacts on the fetus from earlier trimesters in our analysis. Irrespective of these considerations, the sensitivity of the effect to each bombing’s spatial and temporal distance to birth is consistent with the possibility that it reflects exposure to the stressful experience of the bombing.

Previous research in animals using causal study designs has consistently shown that prenatal stress can alter neural development through mechanisms such as disrupted neurotransmitter systems(*16)* and impaired neurogenesis(*15, 16)*. Our findings align with these animal studies, showing that prenatal exposure to air raids during WWII similarly is negatively associated with human neurodevelopment, specifically showing lower gray matter volume (GMV) (reflecting reduced neurogenesis) in corresponding human brain areas, such as frontal cortical areas(*15)*. We also found significant alterations in white matter fiber tracts, such as the corpus callosum, again in line with previously reported animal models(*17)*. However, departing from animal models, we do not find any significant differences in hippocampal areas(*16, 18)*, but rather in the cerebellum and in the striatum, indicating that altered development of the human brain due to prenatal stress deviates in key areas from those animal models. A potential avenue for future research exploring these differences may involve in-vitro human neural models to investigate the underlying mechanisms responsible for alterations in critical areas such as the ventral striatum (a critical hub for reward processing)(*66)* and the cerebellum.

Our results align with smaller prospective human studies indicating that prenatal stress is associated with reduced cortical thickness as well as cognitive and emotional difficulties later in life(*21–23)*. However, our study moves beyond correlational evidence, by leveraging a natural quasi-experiment to shed light on the possible impact of prenatal stress on human fetal neurodevelopment. Our findings are consistent with earlier correlational studies, since we observed an adverse association of prenatal trauma with several frontal brain regions(*21, 22)*, the striatum as well as with the cerebellum(*23)*. Based on previous functional studies implicating prefrontal areas in working memory processes and fluidic IQ(*60, 67, 68)*, and ventral striatal areas in reward processing(*66)*, one may expect that individuals who suffered from GMV reductions in these areas should similarly exhibit reduced working memory capacities (or similarly in fluid IQ)(*69)* or altered primary reward processing. Consistent with this, trauma-related GMV reductions in our sample were indeed associated with lower fluidic IQ and higher BMI (often proposed to reflect altered reward processing)(*70)*. Nonetheless, departing from previous findings, we did not observe any GMV alterations in the temporal lobe(*22, 23)*. While this may be attributed to the limited robustness of previous findings in small sample sizes, it is also possible that previously reported associations in temporal brain areas reflect correlational rather than causal relationships. This highlights the necessity of using larger datasets and - even more importantly - causal inference methods to elucidate the mechanisms underlying altered neurodevelopment due to prenatal stress.

Our results are also generally consistent with previous findings of correlations between prenatal stress and impaired cognitive ability of the child later in life(*23, 26)*. However, our quasi-experimental study revealed a significant indirect effect that was mediated through changes in GMV, with a similar pattern for BMI. Thus, prenatal trauma may affect cognitive ability and obesity due to its influences on neurodevelopment, as measured by Gray Matter Volume (GMV). This suggests that vulnerability to prenatal trauma may be closely interlinked with neurodevelopmental processes that render gray matter development susceptible to detrimental effects of trauma, potentially helping to explain why only some individuals exposed to in-utero stressors exhibit long-term cognitive and health impairments. These individual susceptibility factors may include genetic, epigenetic, or hormonal processes that modulate neurodevelopmental sensitivity(*20, 31, 34, 57)*. For instance, variations in stress hormone regulation (e.g., maternal-fetal cortisol transmission), inflammation, or genetic predispositions affecting neuroplasticity may amplify the impact of prenatal trauma on brain development(*58)*. Identifying such vulnerability factors could be key to understanding differential outcomes and targeting interventions to those most at risk.

Lastly, the fact that our dataset consists solely of UK participants during WWII raises the question whether modern warfare may have different effects in terms of stress and trauma. On the one hand, modern warfare is characterized by precision bombing, which could significantly reduce the effect sizes we observed. On the other hand, mobility may be more severely restricted, as is the case in certain conflicts around the world, so the traumatic effects of war bombings could be substantially magnified, highlighting the need for further investigation into these specific dynamics. Furthermore, although we conducted extensive robustness checks and sensitivity analyses, we cannot fully exclude violations of the as-if-random exposure assumption or residual confounding biases. Nevertheless, viewed alongside converging evidence from mechanistic animal models and prior small-scale human studies, our results are consistent with a direct link between severe prenatal war trauma and altered in-utero brain development.

In conclusion, we report a previously unidentified potential long-term burden of war. Our results underscore the importance of affording special protection to unborn life, and extending evacuations to pregnant women during armed conflicts. The insights gained from our research may have implications for policymakers and international organizations trying to mitigate the problems associated with armed conflicts. Understanding that prenatal exposure to stress can have long-lasting effects on unborn life reinforces the need for urgent and comprehensive policies to protect pregnant women in conflict zones. Interventions could include providing safe havens, stress reduction programs, and psychological support, not only to protect the health of the mother but also to safeguard the developing fetus, potentially reducing the incidence of neurodevelopmental health issues in future generations.

## Acknowledgments

The authors thank Nadja C. Furtner for helpful comments. During the preparation of this work, the authors used OpenAI tools to assist with language editing, grammar, and readability.

## Funding

The research was conducted using UK Biobank resources under application 41060.

The University Research Priority Program ‘Adaptive Brain Circuits in Development and Learning’ (URPP AdaBD) at the University of Zurich (GA, VM, SK, CCR)

NSF Early Career Development Program grant (1942917) (GN)

## Author contributions

Conceptualization: GA, CCR

Methodology: GA, CCR, VM, SK

Investigation: GA, CCR

Visualization: GA, CCR, VM, SK

Funding acquisition: GA, CCR

Project administration: CCR

Supervision: BK, GN, CCR

Writing – original draft: GA, CCR

Writing – review & editing: GA, CCR, RKL, VM, SK, GN, BK

## Data and materials availability

All data, code, and materials used in the analysis can be accessed via the UKBiobank, and data analysis scripts are available on OSF (https://osf.io/vpmhu).

