## Supplementary Materials for "Long-Term Imprint of Prenatal War Trauma on Brain Structure: Evidence from Genetically Informed Neuroimaging"

### Title: War Trauma Impairs Prenatal Brain Development: Evidence from Genetically-Informed Brain Imaging

**The PDF file includes:**

Materials and Methods
Figs. S1 to S4
Tables S1 to S2
References

### Materials and Methods

#### 1. Sample Characteristics and Selection Criteria

We utilized data from the UK Biobank (UKB), which comprises a cohort of 502,617 individuals aged 40 to 69 from across the United Kingdom during data collection(*1*, *2*). UKB participants provided written informed consent, and the North West Multi-Centre Research Ethics Committee approved the study. Participants underwent a series of extensive MR scans, genetic tests, questionnaires, and lifestyle assessments, enabling the exploration of genetic, neural, and behavioral correlations that require large sample sizes to detect(*3*).

Our sample consisted of $N$ = 37,856 individuals who had no missing entries in all relevant variables (MR scans, genotyping data, cognitive, and behavioral phenotypes). Among these, $N$ = 6,368 participants were born between 1939 and 1945. All structural T1 MRI images used in the study underwent automated quality control using the UKB brain imaging processing pipeline (*4*).

#### 2. Measures

##### 2. 1. Air raid exposure and identification strategy

Nazi Germany's terror bombing campaign, starting with the 'Blitz' in 1940, indiscriminately targeted urban centers with the goal of weakening the UK's morale(*5*), but the bombings were also notoriously imprecise. Unlike modern precision strikes, World War II (WWII) bombings had a much larger margin of error, often spanning several kilometers. For example, due to navigational errors, both German and Allied planes mistakenly bombed neutral Switzerland over 70 times, causing many civilian casualties(*5*, *6*). This lack of precision creates a unique opportunity for studying causality in this realm, as it serves as a randomization device. The unpredictability of the air raids, combined with their imprecision, plausibly approximated a random assignment of bombing exposure to unborn children during WWII in the UK, allowing us to examine the effects of prenatal trauma.

To investigate the effect of armed conflicts on the neurodevelopment of unborn children, we used *exposure to air raids* as a proxy for the severity of such prenatal traumatic events. To this end, we utilized data from the British National Archives, which includes meticulously recorded bombing locations from Nazi German air raids, detailing 32,870 air raid locations in the UK between 1939 and 1945 (see Fig. 1B) (*7*).

We then linked the GPS coordinates of those air raids to the time and place of each participant's birth in the UKB sample (Data-Fields 129 & 130; 1km resolution). Specifically, we counted the number of air raids to which a participant in our sample was exposed, considering varying spatial distances (e.g. 2 km, 50 km, 100 km, and 1000 km) and temporal intervals (ranging from 12 months before to 9 months after birth, divided into trimesters). This allowed us to quantify the exposure of each individual to air raids, either before conception, during pregnancy (in utero), or immediately after birth (see Fig. 2).

To make sure the conclusions of our analysis are stable to the choice of a specific spatial radius around the location of the bombings (i.e., <10 km), we conducted a specification curve analysis (see Fig. S1) that estimated how the association between air-raid exposure and grey matter volume (GMV) varied with distance from the birth location. Using our standard controls (birth-location fixed effects, 40 genetic principal components, current living coordinates, sex assigned at birth, birth-year dummies, handedness, and age at scan) and restricting the sample to cohorts born 1939–1945, we ran separate regressions for each radius (e.g., 1–10 km) relating the number of air raids to GMV. Standard errors were clustered within 100 birth-location bins. As expected, the estimated effect sizes decreased monotonically with distance, with distances of 2km to 4km showing the highest significance ($p$ < 3.3×10$^{-5}$). The pattern exhibits a typical effect–variance trade-off: smaller radii capture larger local effects but with lower variance of air raid exposure across subjects within those radii (see Fig S1 Panel C), whereas larger radii pool more events, reducing standard errors but attenuating effects. Specifically, note that this trade off is present in the standard errors of the estimated coefficients, which are inversely proportional to both the square root of the sample size $n$ and the square root of the variance of the explanatory variable, here the # of air raids subjects were exposed to:

$$\mathrm{SE}(\hat{\beta}) \propto \frac{1}{\sqrt{n}} \cdot \frac{1}{\sqrt{\mathrm{Var}(\text{air raid exposure})}} .$$

Specifically, in line with Figure S1, we observe that estimates become more precise when the sample size increases and when air raid exposure shows greater dispersion across individuals. In our data, the drastic increase in the variance of air raid exposure with increasing distance (as shown in Panel C) is mirrored by a corresponding reduction in the standard errors in Panel A. This suggests that much of the gain in precision arises from the increasing variability of the exposure variable rather than from changes in sample size.

Consistent with this, the 1-km estimate had wide standard errors and was not statistically significant. To mitigate this positional uncertainty while retaining locality, we used a 2-km radius in subsequent analyses, which appears the most conservative exposure distance; this also reduces edge misclassification (and any additional uncertainty in raid locations), increases exposed counts, and yields more stable estimates.

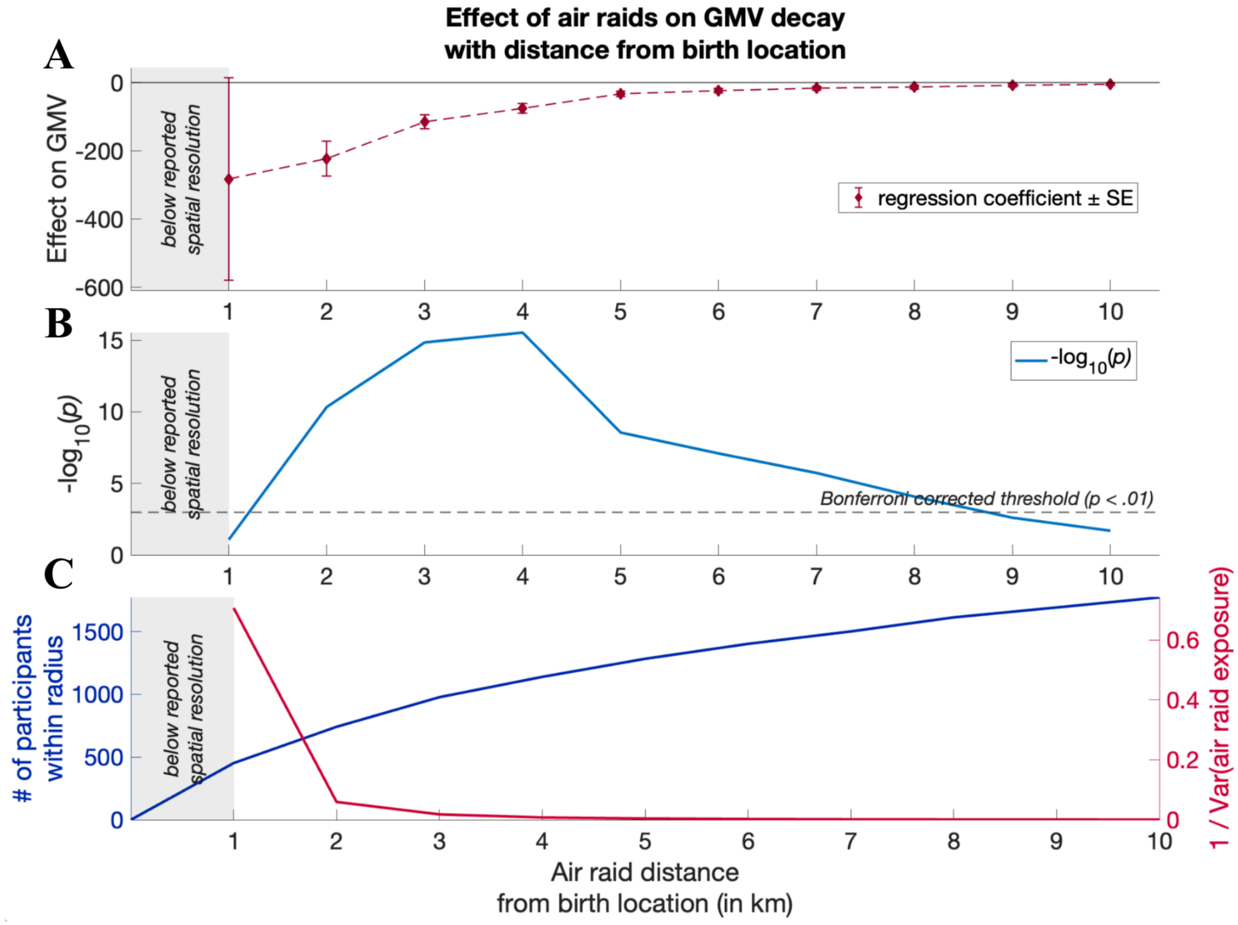


**Fig. S1.** In Panel (A), depicted are regression coefficients ($\beta \pm$ SE) and error bars from ten separate regression models relating the number of air raids in the last trimester to grey matter volume (GMV) across increasing distance bands from birth location. Panel (B) shows the corresponding log-transformed uncorrected *p*-values. The reported resolution of birth locations is 1km (Data-Fields 129 & 130). Each regression uses cohorts born 1939–1945 and includes fixed effects for birth coordinates, 40 genetic principal components, current living coordinates, sex assigned at birth, birth-year dummies, handedness, and age at scan. Standard errors are clustered within 100 birth-location bins. Panel (C) shows the number of participants exposed to air raids in the last trimester (left axis, blue line) across the different radii and the corresponding (inverse) variance in air raid exposure as a function of distance (right axis, red line). The pattern shows a typical effect–variance trade-off: smaller radii capture larger local effects but with less counts of events and with lower dispersion in air raid exposure across participants, whereas larger radii pool more events but attenuating effects. As expected, the negative association is strongest at short distances and attenuates with distance, indicating progressively smaller effects on GMV for exposures farther from the birth location. Distances 2km to 4km show high significance ($Ps < 3.3 \times 10^{-5}$).

Thus, our final spatial analysis considered a 2km radius, and additionally 3 control radii (i.e. 2km, 50km, 100km, 1000km) around the indicated birth location: Air raids within a 2km radius pose a tangible life threat to the expectant mother, as they are very loud, destroy the immediate surroundings, and potentially affect immediate family, neighbors, and close friends. The 50km radius accounts for areas where sirens likely signaled population evacuations during raids, and where likely infrastructure damages that could affect the expectant mother (e.g., food, electricity, and water supply) without direct exposure to bombings. The 100km radius accounts for areas where raids would likely result in sirens-related stress but probably no other direct effects on the expectant mother. Lastly, we included bombings within a 1000 km radius to control for bombings that were reported but where direct raid-related stress should not have occurred. To address potential birth location effects (e.g., being born in East London), we included geographical area dummies and clustered standard errors in 100 location bins (derived from k-means cluster analysis) in all our analyses. This data-driven algorithm identifies naturally occurring clusters by minimizing the distance between birth locations within each group, effectively grouping individuals born in close geographic proximity. This approach avoids bias from arbitrary or manually defined regional boundaries, allowing us to control for local environmental or socioeconomic factors that might otherwise confound the results. This method allows us to systematically control for birth locations and compare different time points (right before or right after birth) for each location separately.

Our temporal analysis examined the three months before estimated conception, the three trimesters of gestation, and the three three-month periods after birth. This allowed us to contrast the exact spatial locations across different times, ensuring comprehensive characterization of the temporal dimension of air raid exposure. As such, bombings within every geographical area (e.g., East London) are evaluated across different time points relative to birth, allowing us to systematically control for self-selection effects that likely occurred during WWII. Specifically, it is nearly impossible to predict in advance for an expectant mother to time the birth of her child, or the location relative to the exact bombing location within her geographical area.

By considering both spatial and temporal dimensions of exposure and controlling for confounding factors, we aim to rigorously examine the potential effects of in-utero trauma on brain structure, contributing to our understanding of the long-term consequences of traumatic events during critical periods of (brain) development.

### 2.2 Geocoding of bombing sites

The British National Archive meticulously recorded 32,870 air raid incidents in the UK between 1939 and 1945 (see Fig. 1B)(*7*). Of those incidents, we identified 9,278 individual bombing locations. We converted these locations into GPS coordinates using google's 'geocoding API'. Entries involving multiple bombing sites, denoted by 'and', '+', and '&' were treated as separate entries. To verify the accuracy of recorded areas, we ensured that the derived GPS coordinates fell within the designated defense regions. Locations that fell outside the indicated defense regions were inspected for spelling errors and other data entry issues (see Supplemental Fig. S2).

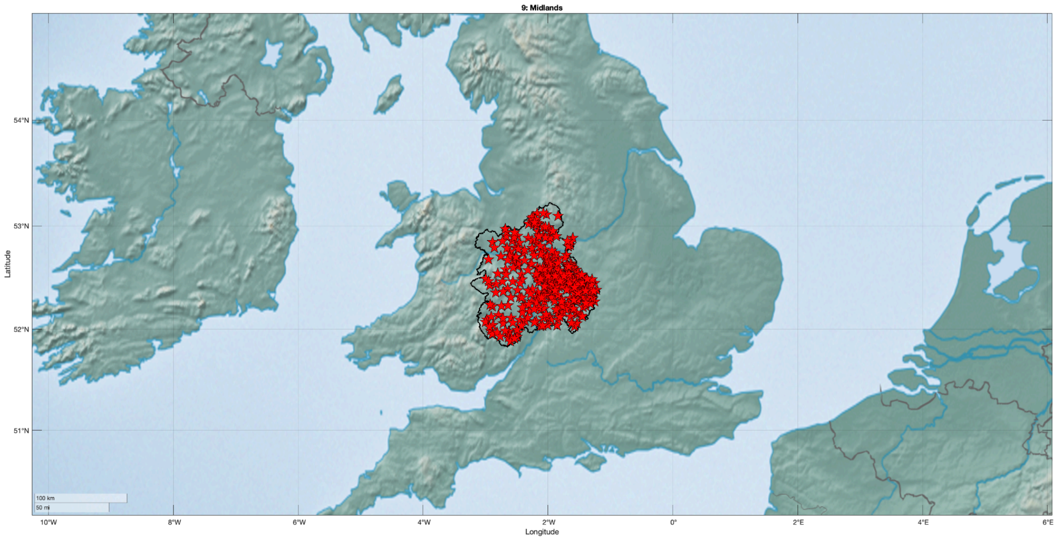


**Fig. S2.** Recorded bombing locations in defense region 9 and its boundaries. To ensure the accuracy of recorded areas, we visually inspected any locations that fell outside of the designated defense region.

We corrected errors in 638 locations where discrepancies arose from the geo-coding software, such as inconsistencies with other information in the original documents (e.g., mismatches with defense regions or other location-related information, obvious spelling errors), or when a location (e.g., a wharf) no longer existed. We determined the most probable coordinates based on historical documents available online when the geo-coding API misinterpreted the nature of the location information, such as confusing a village name with a street name. Excluding locations that were manually corrected from our analyses did not change the results qualitatively.

**2.3 Control Variables**

In all our analyses, we systematically controlled for several genetic, socio-demographic and anthropometric factors, which could potentially confound the observed associations, such as sex, head size and genetic population structure(*8*). Analyses involving MR scans also include controls for age at the time of scan, and standard MR controls such as head scaling and intracranial volume.

Specifically, all of our analyses used the following control variables, which were provided by the UKB:

- Age at the time of brain scan (Data-Field 21003)
- Birth year dummies (Data-Field 33)
- Sex (genetically identified, Data-Field 22001)
- Handedness (Data-Field 1707, categorical variable: Right-handed, Left-handed, ambidextrous, NA)
- The first 40 PCs of the genetic data (Data-Field 22009)
- Volumetric Scaling from T1 head image to standard space (Data-Field 25000)

- Estimated Total intracranial volume (Data-Field 26521)
- *Current* Living Location to control for environmental factors, binned into 100 clusters using *k*-means clustering. (Data-Fields 20074 and 20075)

The description of each Data-Field can be found in the online data showcase of the UKB (http://biobank.ctsu.ox.ac.uk/crystal/search.cgi). The annotated STATA code used to derive the control variables for our analyses can be found on OSF (https://osf.io/vpmhu ).
All variables above were measured on at least one of 3 occasions: (1) the initial assessment visit, (2) the first repeat assessment visit, and (3) the imaging visit. Data from (2) and (3) are only available for a subset of the original sample. In the case where participants provided answers across more than one visit, we used the initial report.

The main analyses are therefore based on the following regression:

$$\begin{aligned}\mathrm{GMV}_i = \beta_0 + \beta_1 \cdot \mathrm{VolumetricScaling}_i + \beta_2 \cdot \mathrm{TotalIntracranialVolume}_i + \sum_{j=1}^{40} \gamma_j \cdot \mathrm{GeneticPC}_{ij} + \delta_1 \cdot \mathrm{AgeAtScan}_i \\ + \delta_2 \cdot \mathrm{SexAtBirth}_i + \delta_3 \cdot \mathrm{BirthYear}_i + \delta_4 \cdot \mathrm{Handedness}_i + \sum_k \theta_k \cdot \mathrm{Bombs}_{ik} \\ + \sum_l \alpha_l \cdot \mathrm{BirthRegion}_{il} + \sum_m \phi_m \cdot \mathrm{LivingRegion}_{im} + \varepsilon_i\end{aligned}$$

The parameters of interest are $\theta_k$, which capture the effect of in-utero exposure to air raids within varying spatial distances (e.g. 2 km, 50 km, 100 km, and 1000 km) and temporal intervals (ranging from 12 months before to 9 months after birth, divided into trimesters). Unless otherwise noted, the primary specification focuses on exposure during the third trimester within 2km radius—the only window that showed a statistically significant association with (total) GMV in the specification-curve analysis (Fig. 2A), while using all other spatial and time frames as controls. In all models, we cluster standard errors by geographic location (100 spatial bins) to accommodate spatially correlated shocks (e.g., areas bombed more frequently) and location-specific unobservables. This yields inference that is robust to within-area correlation and emphasizes comparisons based on temporal variation in exposure among births from the same geographic areas across gestational windows.

#### 2.3.1. Polygenic risk score for fluid intelligence (IQ)

We computed a polygenic risk score (PGS) for fluid intelligence to be used as a control variable for genetic propensity. The procedure initially involved conducting a genome-wide association study (GWAS) on a measure of fluid intelligence (IQ) (Data-Field 20016), constructed by the UKB. The GWAS analyzed a genetically unrelated sample, specifically the non-MRI UKB subsample. Importantly, all participants with (i) MRI data and (ii) their genetic relatives (to the 3rd degree) were excluded from the GWAS. The final GWAS sample size was 186,331. The analysis relied on linear mixed models as implemented in BOLT-LMM v.2.3.6(*9*), and it included control variables for sex, birth year, age at the intelligence assessment, genotyping array and batch, and the first 40 PCs of the genetic data. The outcome was the fluid intelligence score, which was averaged for the subset of respondents with repeated assessment available (to reduce measurement error).

Next, the resulting GWAS summary statistics underwent quality control applied with EasyQC v.9.2.(*10*), which included removal of rare and poorly imputed SNPs, as well as of SNPs deviating more than 0.2 in minor allele frequency from the Haplotype Reference Consortium reference data(*10*, *11*). Finally, the GWAS weights were adjusted for SNP correlations (LD) using the PRS-CS software (*12*). This method restricted the set of SNPs to about 1.1 million quality-controlled HapMap 3 reference SNPs. Finally, the PGI was computed by weighting the UKB genetic data according to their SNP-level weights using PLINK2 software v2.00a3LM (*13*). The final PGS was computed over 1,070,681 SNPs.

#### 2.4 Region of interest (ROI)-level IDPs Processed by the UKB

To examine localized effects in the brain, we used image derived variables (IDPs) provided by the UKB(*4*). These phenotypes include (1) 139 measures of GMV ROIs (Category ID 1101) derived via FSL based on parcellations from the Harvard-Oxford cortical and subcortical atlases and Diedrichsen cerebellar atlas, and (2) 432 measures of white matter fiber integrity derived fromDTI andNODDI (Category ID 134), including fractional anisotropy (FA), mean diffusivity (MD), diffusion tensor mode (MO), axial diffusivity (L1) and radial diffusivity (L2 and L3), intra-cellular volume fraction (ICVF), orientation dispersion index (OD) and isotropic volume fraction (ISOVF).

We regressed each IDP separately on air raid exposure while controlling for the standard control variables listed in 2.3. We corrected for multiple hypothesis testing by adjusting the family-wise-error rate of $\alpha = 0.05$ using permutation tests. For the GMV IDPs, the permutation tests obtained an uncorrected threshold of $P_{uncorr} = 7.7\times10^{-04}$ ($|t_{uncorr}| = 3.363$), and for the white matter fiber IDPs, it resulted in an uncorrected threshold of $P_{uncorr} = 1.8\times10^{-04}$ ($|t_{uncorr}| = 3.7395$).

### 3. Analysis

#### 3.1 Family-Wise-Error Correction using Permutation Tests

To adjust for multiple hypothesis testing, we established a family-wise error-corrected *p*-value threshold using a permutation test method in each analysis. We generated 1,000 datasets with phenotypes randomly permuted (disconnecting the relationship between the dependent and independent variables) (*14*). For each dataset, we calculated regression models for all Imaging Derived Phenotypes (IDPs) and noted the smallest *p*-value from each iteration to form an empirical distribution of the test statistic under the null hypothesis. To determine the family-wise error rate for a specific alpha, we identified the $n^{th}$ = alpha x 1000 smallest *p*-value from the permutation tests as the threshold for uncorrected *p*-values.

#### 3.2 Mediation Analyses

To test whether global GMV differences mediated the association between air raid exposure and fluidic IQ (or BMI), we conducted a mediation analysis in STATA 16. We first performed a structural equation modeling (SEM) analysis to estimate the effect of air raid exposure on fluidic IQ (or BMI) that was mediated via global GMV differences. All SEM equations included the aforementioned standard control variables listed in 2.3.

### 4 Extended Discussion of Results - Possible limitations of the identification strategy

Despite our extensive efforts to identify the causal link between prenatal trauma and brain development, several limitations should be considered when interpreting the results.

First, all participants are from a population sample of adults that were over the age of 40 years at the time of brain imaging. As such, we did not directly measure pregnancy outcomes and immediately followed up on the children in early ages, as common in prospective studies. Despite this limitation, there is no indication that attrition, e.g. through stillbirths during WWII, could have biased our results, as the number of stillbirths even declined during wartime. Similarly, previous research has reported reduced life expectancy in adults exposed to war-related trauma in utero(*15*), which, if present in our sample, may lead to an underestimation of the true extent of trauma-related effects in surviving populations. Nevertheless, we find no indication of potential attrition in our sample.

Furthermore, despite the meticulous efforts of British officials to accurately record all air raid locations, there remains the possibility that some air raids were not documented (see Methods for details on geo-coding). While it is challenging to determine the extent of such a potential bias, the use of geographical area controls should mitigate the impact of such biases, ensuring that our analysis remains robust across different geographical regions.

Further, spatial (self-)selection may have occurred if expectant mothers with lower cognitive ability, possibly due to or associated with socioeconomic constraints, were disproportionately concentrated in heavily bombed urban areas, potentially introducing bias. We have addressed this through spatial controls and via the analysis of effects for different trimesters of pregnancy, which control for the cognitive ability of the mother, by clustering standard errors over 100 locations to isolate the impact of bombings within a given group across different times. Further, our robustness analysis controlling for the polygenic risk for fluid intelligence, a proxy for the expectant mother's cognitive ability, indicates no meaningful change in the effect of bombings. This suggests that the cognitive ability of expectant mothers has not confounded our results

Similarly, temporal self-selection (e.g., delaying family planning based on war prospects) is unlikely, given the unpredictable nature of Nazi Germany's terror bombing campaign. Nevertheless, population statistics from the Office for National Statistics (ONS) showing increased birth rates and robustness analyses including only births during the very first attack wave indicate that the observed effects are not biased by self-selection (i.e., strategic family planning) during the war.

In the following, we discuss those possibilities in greater detail.

*Spatial Self-Selection of Expectant Mothers:* A potential limitation involves the spatial distribution of expectant mothers. Women with lower cognitive ability may have been more likely to live in areas targeted by bombings, such as industrial or urban centers, due to limited employment opportunities or socioeconomic factors. Given that urban areas were strategically significant during World War II, they were heavily bombed. If expectant mothers with lower cognitive ability were overrepresented in these areas, this could introduce bias. Conversely, more educated women, often associated with higher cognitive ability, may have had better access to information about air raid safety and evacuation protocols, enabling them to take protective measures that women with lower cognitive ability may not have fully utilized. To mitigate this bias, we include spatial controls (geographical area dummies and clustered standard errors on over 100 spatial locations), which allows us to compare expectant mothers within the same area across different time points (i.e., before and after the bombing), thus isolating the effect of bombings on neural development. Further, adding the polygenic risk score for cognitive ability as a proxy for parents' cognitive ability did not change our results, indicating that mothers' cognitive ability did not constitute a confounding factor in our analyses.

*Temporal Self-Selection of Mothers:* Another potential concern is that mothers with greater cognitive ability might have been able to avoid exposure to bombings during hot phases of the war, and plan their pregnancy accordingly. However, it is unlikely that prospective mothers could have accurately predicted Nazi Germany's bombing campaign several months in advance, with enough precision to avoid air raids. This is especially true given the chaotic nature of terror bombings, as virtually all of the UK was affected by those air raids. However, by employing spatial and temporal controls, we ensure that even if mothers had been able to temporally avoid bombings by pregnancy planning, our analysis compares mothers from the same spatial location right before and right after exposure to air raids. This helps to account for temporal self-selection and reduces its potential bias.

Further, data from the Office for National Statistics (ONS) in Britain show that fertility rates from 1939 to 1945 rose by approximately 12% compared to pre-war rates, suggesting that family planning remained largely unaffected by the war(*16*). Additionally, stillbirth rates saw a significant decline during the war, dropping from 38.3 stillbirths per 1000 births to 27.6—a nearly 40% reduction(*16*).

*Premature Births*: A further possible concern is that exposure to bombings could lead to an increase in premature births, which might confound the observed effects on fetal development. However, we observed that birth weights—a key indicator of prenatal health—were not significantly affected by bombings, suggesting that premature births are unlikely to drive the results ($p > .1$). This finding is at odds with the possibility that premature birth was a significant confounding factor in this context. This notion is supported by the fact that population-level statistics indicate a decrease in stillbirths(*16*).

*Unmeasured Physical or Social Effects*: While we have systematically controlled for genetic, spatial, and temporal factors, unmeasured physical or social effects related to bombings may not have been fully accounted for. For instance, social support or altered community dynamics in bombed areas could have influenced maternal health in ways that we have not considered. While we cannot fully rule out all possible confounding factors, we minimize the impact of such unobserved variables by incorporating various controls, such as 40 genetic principal components (to account for latent population structure), SES proxied by current residential location,

*Unaccounted Casualties:* Unrecorded casualties, such as stillbirths, may have occurred due to the trauma of air raids. This would result in an underestimation of the true effect size, as the absence of this data would mask the full impact of prenatal exposure to bombings.

*Unaccounted Air Raids:* Some air raids may be missing from the dataset used in our analysis. The omission of these events, as long as it is unsystematic, would similarly lead to an underestimation of the true effect, as exposure levels would be inaccurately reduced. Despite these limitations, our overall conclusions remain robust, though the true impact may be larger than estimated.

### 5. Pre-registration of Analysis Plan and Unplanned Deviations

Prior to data analysis, we pre-registered our analysis plan on the Open Science Framework (OSF, https://osf.io/vpmhu), outlining the operationalization of the dependent variables, control variables, treatment variables, inclusion criteria, and the main and ROI-level analyses.

The main analysis presented in the paper incopproprates slight deviations from this pre-registered plan to better address (unforeseen) confounding factors. Notably, we introduced Total Intracranial Volume (TIV), a well-established standard control variable in neuro-morphometric analysis, as a control variable, to adjust for physiological differences such as head size. We also added current living location (binned into 100 clusters) to control for (present) environmental confounds like socio-economic status. Additionally, we refrained from incorporating birth year - sex interactions into our model due to data sparsity, which would have resulted in too many empty bins, potentially skewing our results.

Finally, we revised our preregistered analysis plan in light of insights from the specification-curve analyses we conducted—analyses that were not available at the time of preregistration—and we accordingly updated our analytis strategy to reflect those findings.

### 6. Code Availability

The software and code used in this study are publicly available, including STATA scripts (OSF, https://osf.io/vpmhu).

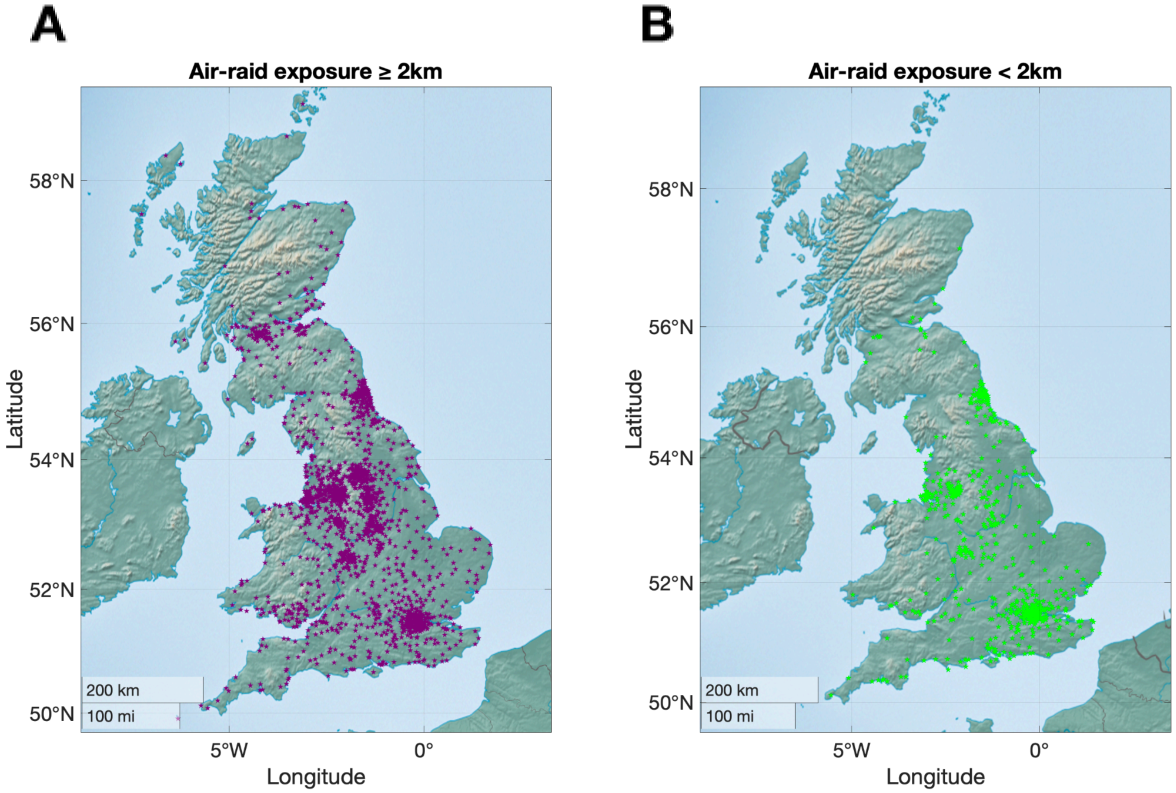


**Fig. S3. |** Both panels show birth locations of 6,664 participants who were in-utero exposed to air raids and had MR scans. 1,539 of them (Panel B) were in-utero exposed to air raids in immediate vicinity (i.e. within 2km radius across all 3 trimesters), while the rest were not directly exposed within 2km radius during pregnancy (Panel A).

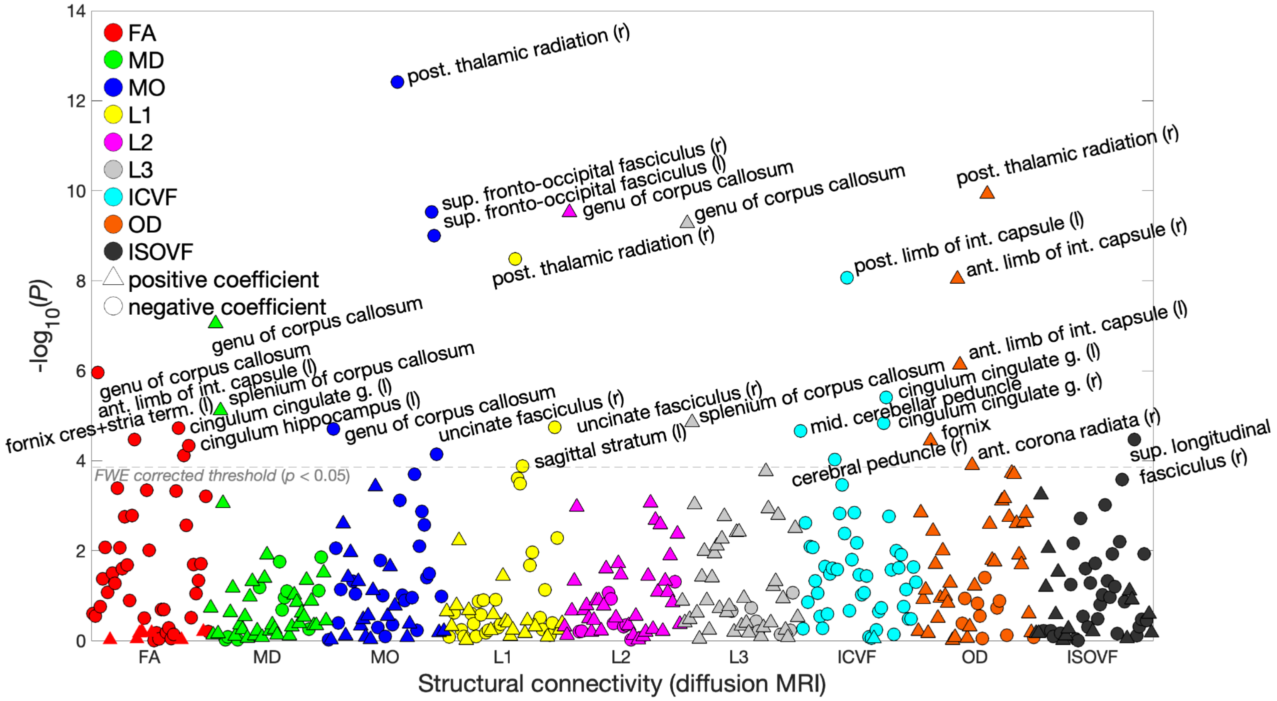


**Fig. S4. |** Analysis of white matter fiber integrity using different DTI and NODDI derived measures (see Methods for Details). Prenatal exposure to air raids (<2km radius in last trimester) resulted in negative effects on white matter fiber integrity, reflected e.g. as lower fractional anisotropy in the *genu of corpus callosum*, lower diffusion tensor mode in the *superior fronto-occipital fasciculus* and in the *posterior thalamic radiation* ($P_{FWE} < .05$), all crucial white matter structures related to important cognitive abilities (e.g. short-term memory and fluidic IQ)(*17*–*19*).

**Table S1.** | Regression coefficients of all regions of interest (ROIs) whose gray matter volume (GMV) demonstrates a significant association with air raid exposure within 2 km radius during the last trimester, as illustrated in Figure 3B. These ROIs have passed the family-wise error correction with an uncorrected threshold of $P_{uncorr}$ = 7.7×$10^{-04}$ ($|t_{uncorr}|$ = 3.363).

| **Gray Matter Volume (GMV) ROI** | **beta** | **SE** | $p_{uncorr}$ | $t_{uncorr}$ |
|---|---|---|---|---|
| Frontal Pole (l) | -0.029 | 0.008 | 0.000 | -3.887 |
| Inferior Frontal Gyrus, pars opercularis (r) | -0.034 | 0.007 | 0.000 | -4.673 |
| Precentral Gyrus (r) | -0.043 | 0.011 | 0.000 | -4.072 |
| Frontal Medial Cortex (l) | -0.033 | 0.009 | 0.000 | -3.872 |
| Frontal Medial Cortex (r) | -0.047 | 0.010 | 0.000 | -4.606 |
| Juxtapositional Lobule Cortex (formerly Supplementary Motor Cortex) (l) | -0.035 | 0.009 | 0.000 | -4.028 |
| Supramarginal Gyrus, posterior division (l) | -0.045 | 0.010 | 0.000 | -4.590 |
| Putamen (r) | -0.031 | 0.008 | 0.000 | -3.907 |
| Amygdala (r) | -0.042 | 0.008 | 0.000 | -5.244 |
| Ventral Striatum (l) | -0.042 | 0.009 | 0.000 | -4.722 |
| Ventral Striatum (r) | -0.051 | 0.010 | 0.000 | -4.904 |
| V Cerebellum (l) | -0.045 | 0.012 | 0.000 | -3.880 |
| VI Cerebellum (l) | -0.033 | 0.009 | 0.000 | -3.711 |
| VI Cerebellum (r) | -0.038 | 0.010 | 0.000 | -3.950 |
| Crus II Cerebellum (l) | -0.055 | 0.014 | 0.000 | -4.057 |
| VIIb Cerebellum (l) | -0.045 | 0.007 | 0.000 | -6.251 |
| VIIb Cerebellum (r) | -0.044 | 0.011 | 0.000 | -4.075 |
| VIIIa Cerebellum (l) | -0.046 | 0.007 | 0.000 | -6.403 |
| VIIIa Cerebellum (r) | -0.050 | 0.008 | 0.000 | -6.355 |
| Vermis IX Cerebellum | -0.048 | 0.009 | 0.000 | -5.298 |
| IX Cerebellum (r) | -0.044 | 0.011 | 0.000 | -3.869 |

**Table S2.** | Regression coefficients of all regions of interest (ROIs) whose white matter fiber integrity measures (DTI and NODDI based) showed significant associations with air raid exposure within 2 km radius during the last trimester, as illustrated in Figure S4. These ROIs have passed the family-wise error correction with an uncorrected threshold of $P_{uncorr} = 1.8\times10^{-04}$ ($|t_{uncorr}| = 3.7395$).

| **White Matter Fiber ROI** | **beta** | **SE** | $p_{uncorr}$ | $t_{uncorr}$ |
|---|---|---|---|---|
| FA in genu of corpus callosum on FA skeleton | -0.051 | 0.010 | 0.000 | -4.873 |
| FA in ant. limb of int. capsule on FA skeleton (l) | -0.056 | 0.013 | 0.000 | -4.148 |
| FA in cingulum cingulate g. on FA skeleton (l) | -0.034 | 0.008 | 0.000 | -4.277 |
| FA in cingulum hippocampus on FA skeleton (l) | -0.031 | 0.008 | 0.000 | -3.955 |
| FA in fornix cres+stria term. on FA skeleton (l) | -0.049 | 0.012 | 0.000 | -4.072 |
| MD in genu of corpus callosum on FA skeleton | 0.044 | 0.008 | 0.000 | 5.349 |
| MD in splenium of corpus callosum on FA skeleton | 0.031 | 0.007 | 0.000 | 4.475 |
| MO in genu of corpus callosum on FA skeleton | -0.030 | 0.007 | 0.000 | -4.269 |
| MO in post. thalamic radiation on FA skeleton (r) | -0.040 | 0.005 | 0.000 | -7.262 |
| MO in sup. fronto-occipital fasciculus on FA skeleton (r) | -0.047 | 0.007 | 0.000 | -6.302 |
| MO in sup. fronto-occipital fasciculus on FA skeleton (l) | -0.073 | 0.012 | 0.000 | -6.109 |
| MO in uncinate fasciculus on FA skeleton (r) | -0.040 | 0.010 | 0.000 | -3.968 |
| L1 in post. thalamic radiation on FA skeleton (r) | -0.041 | 0.007 | 0.000 | -5.916 |
| L1 in sagittal stratum on FA skeleton (l) | -0.032 | 0.009 | 0.000 | -3.822 |
| L1 in uncinate fasciculus on FA skeleton (r) | -0.044 | 0.010 | 0.000 | -4.290 |
| L2 in genu of corpus callosum on FA skeleton | 0.054 | 0.009 | 0.000 | 6.297 |
| L3 in genu of corpus callosum on FA skeleton | 0.050 | 0.008 | 0.000 | 6.210 |
| L3 in splenium of corpus callosum on FA skeleton | 0.033 | 0.008 | 0.000 | 4.342 |
| ICVF in mid. cerebellar peduncle on FA skeleton | -0.039 | 0.009 | 0.000 | -4.248 |
| ICVF in cerebral peduncle on FA skeleton (r) | -0.041 | 0.011 | 0.000 | -3.904 |
| ICVF in post. limb of int. capsule on FA skeleton (l) | -0.042 | 0.007 | 0.000 | -5.756 |
| ICVF in cingulum cingulate g. on FA skeleton (r) | -0.044 | 0.010 | 0.000 | -4.335 |
| ICVF in cingulum cingulate g. on FA skeleton (l) | -0.041 | 0.009 | 0.000 | -4.615 |
| OD in fornix on FA skeleton | 0.037 | 0.009 | 0.000 | 4.134 |
| OD in ant. limb of int. capsule on FA skeleton (r) | 0.029 | 0.005 | 0.000 | 5.745 |
| OD in ant. limb of int. capsule on FA skeleton (l) | 0.030 | 0.006 | 0.000 | 4.952 |
| OD in ant. corona radiata on FA skeleton (r) | 0.034 | 0.009 | 0.000 | 3.833 |
| OD in post. thalamic radiation on FA skeleton (r) | 0.041 | 0.006 | 0.000 | 6.441 |
| ISOVF in sup. longitudinal fasciculus on FA skeleton (r) | -0.034 | 0.008 | 0.000 | -4.148 |
